# *Streptomyces* volatile compounds influence exploration and microbial community dynamics by altering iron availability

**DOI:** 10.1101/396606

**Authors:** Stephanie E. Jones, Christine A. Pham, Joseph McKillip, Matthew Zambri, Erin E. Carlson, Marie A. Elliot

**Affiliations:** Department of Biology and Michael G. DeGroote Institute for Infectious Disease Research, McMaster University, Hamilton, Ontario, Canada; Department of Chemistry, University of Minnesota, Minneapolis, Minnesota, USA

## Abstract

Bacteria and fungi produce a wide array of volatile organic compounds (VOCs), and these can act as infochemicals or as competitive tools. Recent work has shown that the VOC trimethylamine (TMA) can promote a new form of *Streptomyces* growth, termed ‘exploration’. Here, we report that TMA also serves to alter nutrient availability in the area surrounding exploring cultures: TMA dramatically increases the environmental pH, and in doing so, reduces iron availability. This, in turn, compromised the growth of other soil bacteria and fungi. In contrast, *Streptomyces* thrives in these iron-depleted niches by secreting a suite of differentially modified siderophores, and by upregulating genes associated with siderophore uptake. Further reducing iron levels by siderophore piracy, limiting siderophore uptake, or growing cultures in the presence of iron chelators, unexpectedly enhanced exploration. Our work reveals a new role for VOCs in modulating iron levels in the environment, and implies a critical role for VOCs in modulating the behaviour of microbes and the makeup of their communities.

## INTRODUCTION

Bacteria and fungi frequently live in densely populated multispecies communities. These microbes produce a vast array of molecules capable of modulating community dynamics, including specialized metabolites and volatile organic compounds (VOCs) (1). Soil environments are particularly complex: not only are they home to multitudes of microbes, but they are also heterogeneous systems, containing solid microenvironments and nutrient gradients connected by networks of water- and air-filled pores (2). To date, the majority of studies on interspecies competition in microbial communities have focused on the effects of specialized metabolites. These compounds effectively mediate microbial communication and competition, but their effects are limited to interactions occurring in close proximity, due to their limited diffusion capabilities. In contrast, VOCs are low-molecular-weight compounds that are capable of rapidly diffusing across water channels and air pockets, and consequently, can act as longer-range signals (3). The biological roles of VOCs are only now starting to be dissected, and initial studies are showing that they can have broad effects on both their producing organisms and their neighbours. Indeed, VOCs can alter the antibiotic resistance profiles of bacteria, act as antifungal or antibiotic compounds, promote group behaviours such as motility and biofilm formation, and induce widespread changes in the gene expression of nearby microbes (4).

One group of prolific volatile producers are the *Streptomyces* bacteria (5–7). In the soil, these bacteria are best known for their ability to produce a vast array of specialized metabolites, and for their complex, filamentous life cycle (8, 9). Recent work has, however, revealed that *Streptomyces* species also use volatile compounds to promote an alternative growth strategy known as ‘exploration’ (10,11). In the model species *Streptomyces venezuelae*, exploration is initiated in response to the production of the VOC trimethylamine (TMA), which promotes the rapid and unrelenting spreading of explorer cells across surfaces. TMA production dramatically alters the surrounding environment, raising the pH to levels approaching 9.5, and further serves as a *Streptomyces* communication signal, inducing exploration in physically separated streptomycete colonies. TMA-mediated induction of exploration appears to be a function of its alkalinity, as other alkaline VOCs (*e.g.* ammonia) can induce exploration in a similar manner. TMA is also effective as a weapon against non-streptomycetes: both exploring *Streptomyces* colonies and TMA solutions reduce the survival of other soil bacteria, including *Bacillus subtilis* and *Micrococcus luteus* (11).

The effects of TMA on environmental alkalinization, *Streptomyces* exploration, and the growth of other microbes suggest far-reaching effects for this VOC. How TMA affects microbial community dynamics and impacts the growth of other microbes is, however, not clear. Here, we demonstrate that TMA emitted by *Streptomyces* explorer cells reduces the survival of other soil bacteria and fungi by starving them of iron – a micronutrient that is critical for microbial growth and viability. Within these self-induced iron-depleted environments, *Streptomyces* thrive by secreting siderophores and rewiring gene expression to maximize siderophore uptake. We show that iron depletion by other microbes, or by iron chelators, can enhance *Streptomyces* exploration, revealing that low iron is a driver of exploratory growth. Taken together, our results reveal a new way in which *Streptomyces* can alter the availability of environmental iron, and in turn influence the growth and behavior of themselves and other members of the surrounding microbial communities. Our findings further suggest that iron depletion can activate a positive feedback loop that contributes to the relentless expansion of exploring cultures.

## MATERIALS AND METHODS

### Strains, plasmids, media and culture conditions

Strains, plasmids and primers used in this study are listed in **Table 2**. *S. venezuelae* ATCC 10712 was grown on MYM (maltose, yeast extract, malt extract) agar for spore stock generation, and for examining the behavior of classically developing cultures. Exploration was investigated on YP (yeast extract, peptone) agar or in association with yeast on YP agar supplemented with dextrose/glucose (YPD). Non-exploring controls were grown by themselves on YPD agar. For iron experiments, plates were supplemented with the indicated concentration of 2,2′-dipyridyl (0-360 µM) or FeCl_3_ (0-10 mM). All strains were grown at 30°C, except those experiments involving TMA, which were conducted at room temperature in a fume hood. Prior to growing on plates, *S. venezuelae* was grown in liquid MYM at 30°C, and 5 µL were spotted to agar plates. *Amycolatopsis* strains were also grown in liquid MYM at 30°C, and 5 µL of the overnight culture was spotted alone or directly beside *S. venezuelae* on the surface of YP agar medium, or *S. coelicolor* M145 on the surface of R2YE agar medium. *S. coelicolor* was spotted directly from a spore stock. All plates were incubated for up to 14 days. Colony surface areas were measured using ImageJ (12). *E. coli* strains were grown in or on LB (Lysogeny Broth) medium or in SOB (super optimal broth) medium. DH5α and ET12567/pUZ8002 strains were grown at 37°C, and BW25113/pIJ790 was grown at 30°C or 37°C.

**Table 1.** *S. venezuelae* siderophore production and uptake genes

| Genes |  |
| --- | --- |
| **Synthesis clusters** | **Predicted siderophore** |
| *sven_0503-17* | Thiazostatin |
| *sven_2566-77* | Desferrioxamine |
| *sven_5413-26* | Unknown siderophore |
| *sven_5472-82* | Rhizobactin-like cyptic siderophore |
| *sven_7032-80* | Coelichelin |
| **Uptake systems** | **Annotation** |
| ***bldK* homologs** |  |
| *sven_4759-63* |  |
| *sven_5150-54* |  |
| *sven_4765-69* |  |
| *sven_4820-23* |  |
| *sven_5369-73* |  |
| *sven_6981-85* |  |
| *sven_7098-02* |  |
| *sven_7135-39* |  |
| *sven_7154-58* |  |
| ***other*** |  |
| *sven_0776-78* | Salmycin transporter CdtABC |
| *sven_0164-66* | Putative siderophore transporter system |
| *sven_1997* | Putative iron-siderophore uptake system |
| *sven_1954-57* | Ferrous iron transport system EfeUOB |

**Table 2.** Bacterial strains and plasmids

| Strain | Genotype, description, or use | Reference or source |
| --- | --- | --- |
| ***Streptomyces*** |  |  |
| *Streptomyces venezuelae* ATTC 10712 |  | Gift from M. Buttner |
| E329 | ATTC 10712 ΔSVEN_4759::[aac(3)IV-oriT] | This work |
| E330 | ATTC 10712 ΔSVEN_5151::TOPO 2.1 | This work |
| E330 | ATTC 10712 ΔSVEN_4759::[aac(3)IV-oriT] + ΔSVEN_5151::TOPO 2.1 | This work |
| E331 | ATTC 10712 ΔSVEN_2570-73::[aac(3)IV-oriT] | This work |
| *Streptomyces coelicolor* M145 |  | (38) |
| ***Amycolatopsis*** |  |  |
| *Amycolatopsis orientalis*<br>*sp. lurida* |  | Gift from G. Wright |
| ***E. coli*** |  |  |
| DH5α | Plasmid construction and general subcloning | Invitrogen |
| ET12567/pUZ8002 | Generation of methylation-free plasmid DNA and conjugation into *Streptomyces* | (39, 40) |
| BW25113/pIJ790 | Mutations in cosmid DNA | (15) |
| **Plasmids or cosmids** |  |  |
| 3D02 | *S. venezuelae* cosmid carrying *SVEN_4759* | Gift from M. Bibb and M. Buttner |
| 5-E05 | *S. venezuelae* cosmid carrying *SVEN_2570-73* | Gift from M. Bibb and M. Buttner |
| TOPO 2.1 | Plasmid used for disruption of *SVEN_5151* | Invitrogen |
| pIJ790 | Temperature-sensitive plasmid carrying λ-RED genes | (15) |
| pIJ773 | Plasmid carrying the apramycin knockout cassette | (15) |
| pIJ10701 | Plasmid carrying the *hyg-oriT* cassette | (15) |
| pIJ780 | Plasmid carrying the *vio-oriT* cassette | (15) |

For iron growth assays, *B. subtilis* and *M. luteus* were grown in LB medium, while *S. cerevisiae* was grown in YPD medium. Each strain was grown in 10 mL liquid media shaking overnight at 30°C. To quantify strain survival in response to iron chelation on solid medium, each strain was subcultured and grown to an OD_600_ of 0.8, before being diluted 1/5000, with 100 µL being spread on agar medium containing 0-360 μM 2,2’ dipyridyl (hereafter referred to as dipyridyl). Colony numbers were quantified after 2 days. To quantify the survival of each strain in response to iron chelation in liquid medium, varying amounts of overnight cultures of each strain were added to 1.5 mL fresh media to give an OD_600_ of 0.1 in 48-well plates. Plates were shaken at 30°C in a plate reader and OD600 readings were taken every 30 min for 8 hours.

### Identification and analysis of desferrioxamines

To analyze metabolite production by explorer cultures, agar from each culture plate (alongside a medium control) was diced, placed in a 50 mL falcon tube, and frozen at -80°C. The cultures/agar were then dried by lyophilization. Extraction of metabolites was performed by adding 45 mL of 50:50 n-butanol/ethyl acetate to each falcon tube and rotating overnight at room temperature. The extracts were filtered, and the solvent removed under vacuum at room temperature (Genevac EZ-2 Series Personal Evaporator, method low + medium BP, lamp off).

All mass spectrometry experiments were conducted using UPLC-ESI-QTOF MS instrumentation (Agilent, 6540), as described previously (13, 14). Each extract was dissolved in 400 µL of 50:50 H_2_O/acetonitrile (Fisher, LC-MS grade), of which 2 µL were injected onto a C18 column (Agilent, Zorbax, 2.1 x 50 mm, 1.8 µm). Compounds were separated using a constant flow of 0.4 mL/min and the following gradient: 0-3 min at 0% B (A, 95:5:0.1%, H_2_O/acetonitrile(ACN)/formic acid and B, 95:5:0.1% ACN/H_2_O/formic acid), 3-17 min at 0-100% B, and 17-20 min at 100% B. Accurate mass data were acquired in triplicate in both profile and centroid mode with source/fragmentor voltage of 100 V, positive mode ion detection between 100–1,700 *m/z*, gas temperature of 325°C, and capillary voltage of 3,500 V. Fragmentation data were acquired with a fixed collision energy of 35 V with positive mode product ion detection between 50-1,650 *m/z*. Data were processed using Agilent Masshunter Qualitative Analysis and Origin.

### Construction of deletion strains and mutant complementation

In-frame deletions of *sven_4759* (*bldK* homolog) and *sven_2570-73* (*desA-D*) were generated using ReDirect technology (Gust et al., 2003). For each of *sven_4759* and *desA-D*, the coding sequence (from start to stop codon) was replaced with an *oriT*-containing apramycin resistance cassette. For mutation of *sven_5151* (a second *bldK* homolog), the gene was disrupted in the chromosome. A 1,406-bp region of the gene was amplified and cloned into the TOPO vector (Invitrogen). Mutant cosmids/disruption plasmids were introduced into the non-methylating *E. coli* strain ET12567/pUZ8002 prior to conjugation into wild type *S. venezuelae*. For creating the Δ*sven_4759*Δ*sven_5151* double mutant strain, the ET12567/pUZ8002 strain carrying the *sven_5151* TOPO construct was introduced into the Δ*sven_4759*:apramycin strain. Resulting exconjugants were screened for double-crossover recombinants (in the case *sven_4759 bldK* and *sven_2570-73*) or single-crossover integration (in the case of *sven_5151* and Δ*sven_4759*Δ*sven_5151*). Correct replacement of *sven_4759* or *sven_2570-73*, or disruption of the *sven_5151* coding sequence was confirmed using diagnostic PCR combinations (see **Table 3**). The Δ*sven_4759*Δ*sven_5151* double mutant phenotype was complemented using a cosmid carrying the wild type *sven_4759* sequence, along with the downstream cluster to account for any polar effects. To enable effective selection for cosmid integration in the *S. venezuelae* chromosome, the ampicillin resistance gene on the cosmid backbone was replaced with an *oriT*-containing viomycin resistance cassette, using the ReDirect protocol (15).

**Table 3.**
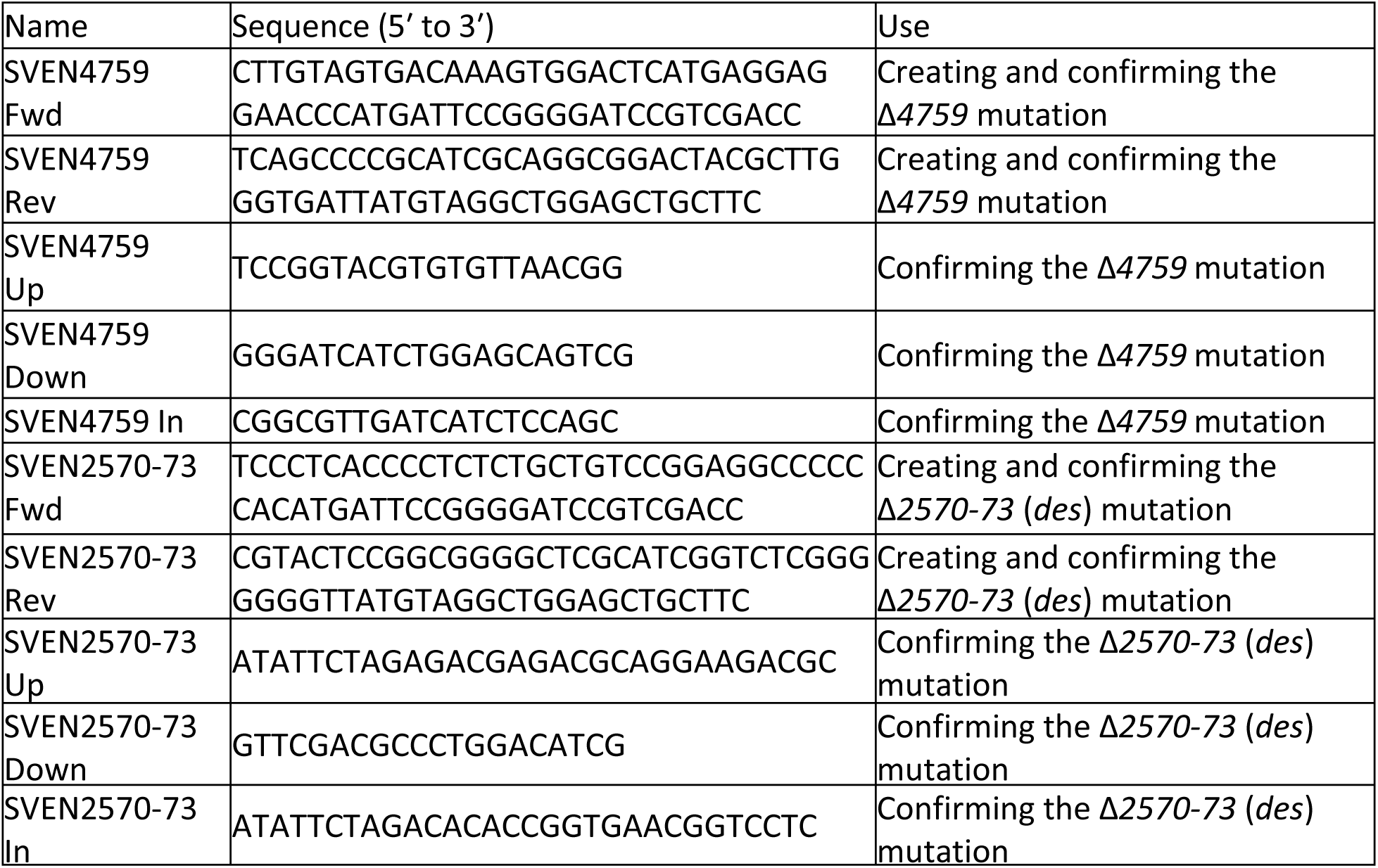
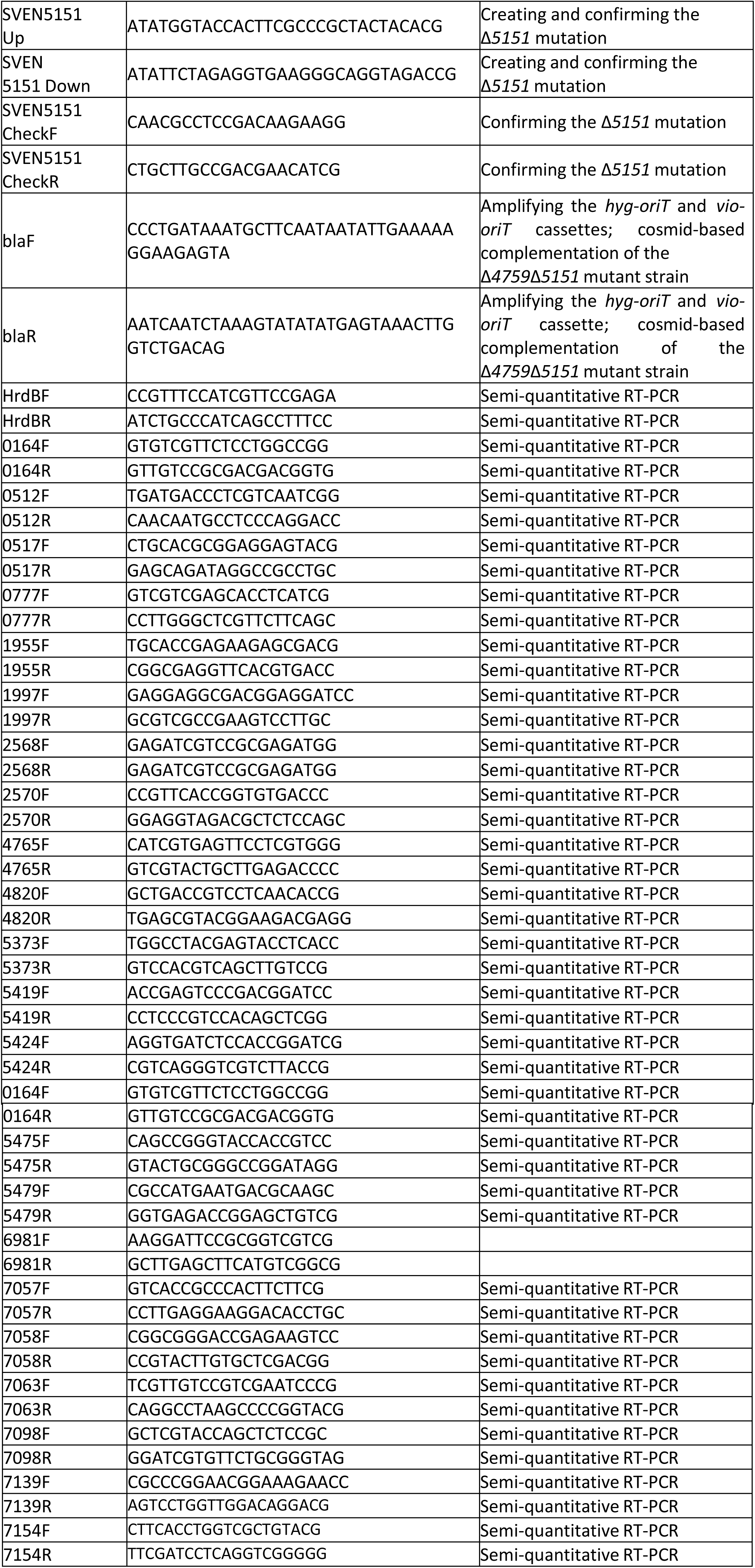
Oligonucleotides used in this study

### RNA isolation and RT-PCR analysis

RNA was isolated as described previously from two replicates from each of wild type and Δ*sven_4759*Δ*sven_5151 S. venezuelae* exploring colonies grown for 8 days on YP agar plates (we were unable to isolate high quality RNA from exploring *S. venezuelae* at later time points). For all replicates, contaminating DNA was removed using Turbo DNase (Life Technologies), and was confirmed to be DNA-free by PCR. RNA quality and purity were assessed using a Nanodrop spectrophotometer. RNA integrity was further analyzed by agarose gel electrophoresis prior to reverse transcription-PCR (RT-PCR) analysis.

One microgram of RNA was used as template for reverse transcription using gene-specific primers (see **Table 3**) and Superscript III polymerase (Invitrogen), according to the manufacturer’s instructions. The resulting cDNA then served as template for PCR amplification using Taq DNA polymerase and the gene-specific primers listed in **Table 3**. The number of cycles was optimized to ensure products were detected in the linear range of amplification. Negative controls containing nuclease-free water instead of reverse transcriptase were included to ensure RNA samples and other reagents did not contain residual contaminating DNA (‘no RT’ control). cDNA corresponding to the gene encoding the vegetative sigma factor *hrdB* was amplified as a positive control for RNA levels and RNA integrity. Ten microliters of each PCR were separated on a 1.5% agarose gel and visualized by staining with ethidium bromide. All reactions were conducted in triplicate, using two independently isolated RNA samples.

### Assays for volatile-mediated phenotypes

To quantify how *S. venezuelae* VOCs affected the survival of other bacteria or fungi, *S. venezuelae* was grown in a small petri dish containing YP or YPD agar. The small dish was placed inside a larger dish containing agar or agar supplemented with 1 mM FeCl_3_. LB agar was used for the growth of *B. subtilis* and *M. luteus*, while YPD agar was used for *S. cerevisiae*. *S. venezuelae-*inoculated plates were grown for 10 days, after which the indicator organisms were spread on the surrounding plates. *B. subtilis, M. luteus*, or *S. cerevisiae* were subcultured and grown to OD_600_ of 0.8 in liquid LB/YPD, after which the cultures were diluted 1/10,000 and 50 µL were then spread on the agar. Colony numbers on the outer plate were then quantified after 48 hours.

Measuring how iron supplementation affected the responses of microbes to TMA involved adding either 1.5 mL of commercially available TMA solutions (Sigma) diluted to 0.9 w/v, or water (negative control) to sterile plastic containers. These were then placed in a petri dish containing 50 mL LB or YPD agar, with or without 1 mM FeCl_3_ supplementation. *B. subtilis, M. luteus*, or *S. cerevisiae* were subcultured and grown to an OD_600_ of 0.8 in liquid LB/YPD before 100 μL of the subculture were spread on each plate. Plates were incubated at room temperature for 48 hours, before cells were scraped into tubes containing 2 mL liquid LB or YPD and vigorously mixed. Dilution series’ were used to measure the OD_600_ of the resulting cell suspensions. A minimum of four biological replicates were assessed, alongside two technical replicates in each instance.

## RESULTS

### Environmental iron availability impacts the survival of bacteria and fungi

Iron is an essential nutrient for most organisms; however, its acquisition is a major challenge. Cells use ferrous iron (Fe^2+^), however, iron in the environment exists predominantly in its poorly soluble ferric form (Fe^3+^). To facilitate iron acquisition, bacteria release iron-chelating siderophores (16). These small molecules bind ferric iron, and the resulting siderophore-iron complexes are taken back up into cells through dedicated membrane transporters, after which iron is released in its ferrous form. In alkaline environments, iron solubility drops by ∼1000 fold with each unit rise in pH, due to ferric iron forming stable complexes with hydroxide ions. This further lowers iron solubility and reduces iron binding by siderophores (17–19).

*S. venezuelae* exploration requires an alkaline environment, which they create by releasing the volatile compound TMA (**Fig. S1**). TMA emission also results in dramatically decreased survival of other soil-dwelling bacteria (11). Consequently, we wondered whether low iron availability could explain the reduced survival observed for other microbes exposed to exploration-associated VOCs. To address this possibility, we first tested the extent to which low iron affected the growth of the soil bacteria *B. subtilis* and *M. luteus*, as well as the fungus *Saccharomyces cerevisiae*.

We first compared the growth of these strains on solid medium, relative to medium supplemented with the iron-specific chelator 2,2′-dipyridyl (hereafter referred to as dipyridyl) (160 or 320 µM). For *M. luteus*, colony numbers were equivalent on LB and on LB with 160 µM dipyridyl, although colonies were smaller when growing on dipyridyl, suggesting that low iron slowed the growth of these organisms. With 320 µM dipyridyl, however, no *M. luteus* colonies survived (**Fig. 1A**). In contrast, for *B. subtilis* we observed a linear growth decrease on LB medium containing dipyridyl compared with LB medium alone: on 160 µM dipyridyl, colony numbers were reduced by ∼30%, and they dropped a further 30% on 320 µM dipyridyl (**Fig. 1A**). Finally, in the case of *S. cerevisiae*, colony numbers decreased by an average of ∼40% on YPD with 160 µM dipyridyl; on plates containing 320 µM dipyridyl, colonies failed to grow altogether (**Fig. 1A**).

**Fig. 1.**
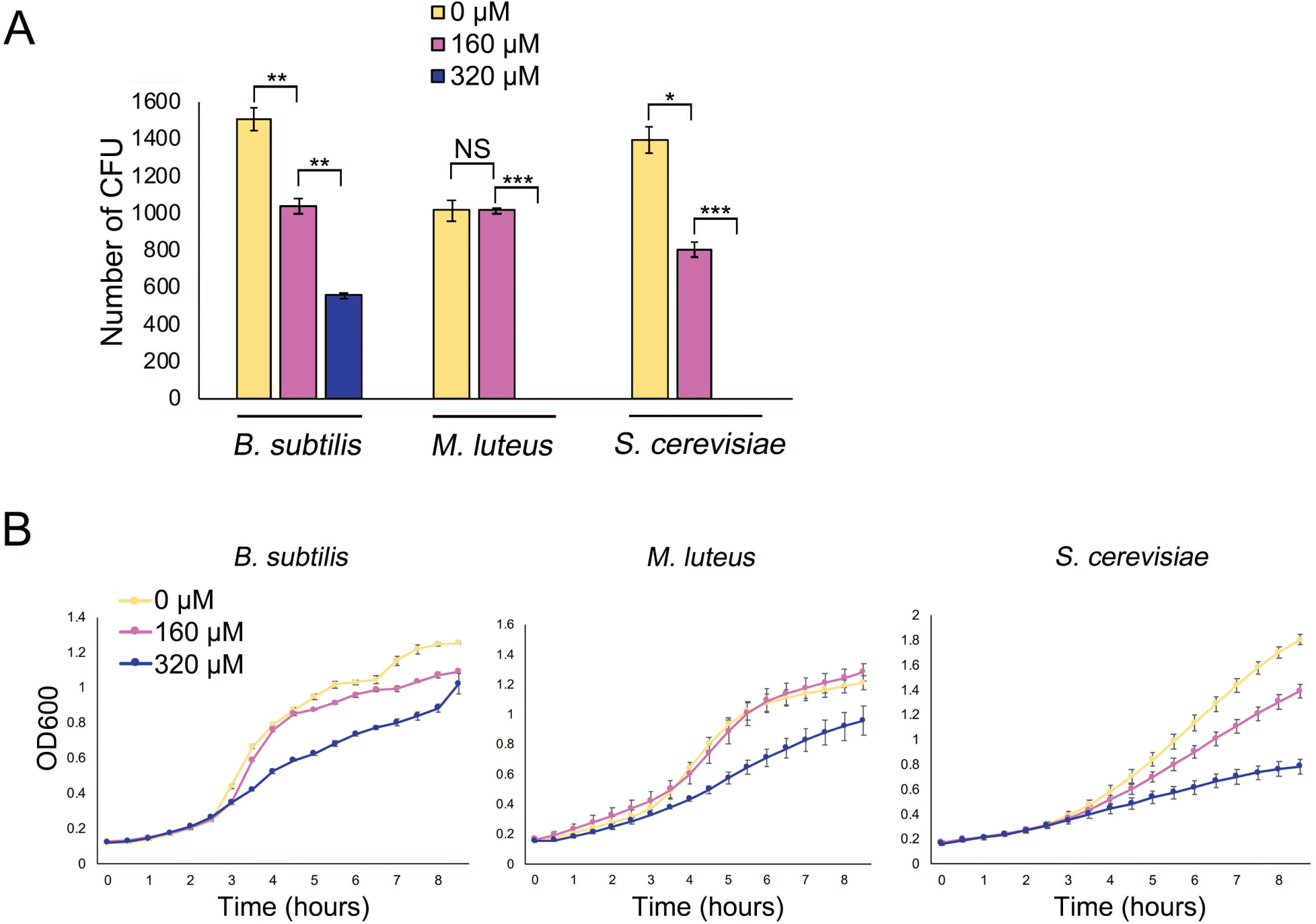
Iron availability impacts the growth and survival of bacteria and fungi. **A.** Quantification of colony forming units (CFU) for *B. subtilis* and *M. luteus* on LB agar medium and *S. cerevisiae* on YPD agar medium, supplemented with 0, 160 or 320 µM 2,2′-dipyridyl. Plates were incubated for 48 h. All values represent the mean ± standard error for three or four biological replicates. Asterisks indicate statistically significant differences (*: *p*-values of 0.05 to 0.01; **: *p*-values of 0.01 to 0.005; ***: *p*-values below 0.005, NS: no significant difference) as determined by a student’s t-test. Note that *M. luteus* and *S. cerevisiae* did not grow at 320 µM 2,2′-dipyridyl. **B.** Growth curves of *B. subtilis* and *M. luteus* in liquid LB medium and *S. cerevisiae* in liquid YPD medium, supplemented with 0, 160 or 320 µM 2,2′-dipyridyl and grown for 8 h. Values represent the mean ± standard error for three biological replicates.

We also tested the effect of dipyridyl on the liquid culture growth of each of these organisms. Each microbe was grown in liquid medium (LB for *B. subtilis* and *M. luteus*; YPD for *S. cerevisiae*), and growth was compared with that in medium supplemented with 160 µM or 320 µM dipyridyl. As we saw for the solid-grown cultures, the growth rate for each strain decreased as dipyridyl concentrations increased (**Fig. 1B**). Collectively, these experiments verified that iron was important for the growth of these microorganisms.

### Iron supplementation rescues microbial growth in the presence of explorer cells

Having demonstrated that sufficient iron was essential for robust growth by *B. subtilis, M. luteus* and *S. cerevisiae*, we sought to test our hypothesis that the volatile compounds produced by exploring *S. venezuelae* reduced the survival of other microbes by creating an alkaline, iron-deficient environment. We set up a small petri dish of YPD agar (non-exploring medium) or YP agar (exploring medium) inside a larger dish of solid media, either alone or supplemented with additional iron. The small petri plates were inoculated with *S. venezuelae* and incubated for 10 days, after which *B. subtilis, M. luteus* or *S. cerevisiae* were spread on the larger, surrounding agar plates (**Fig. 2A**). Growth of each indicator microbe was assessed after 2 days. When grown adjacent to either YP plates without *Streptomyces* inoculum or to non-exploring *S. venezuelae* cultures on YPD medium, colony numbers for each microbe were similar on all plates, irrespective of iron supplementation (**Fig. 2B**). As extra iron did not enhance growth, it suggested that iron was unlikely to be limiting for the growth of these organisms under these conditions.

**Fig. 2.**
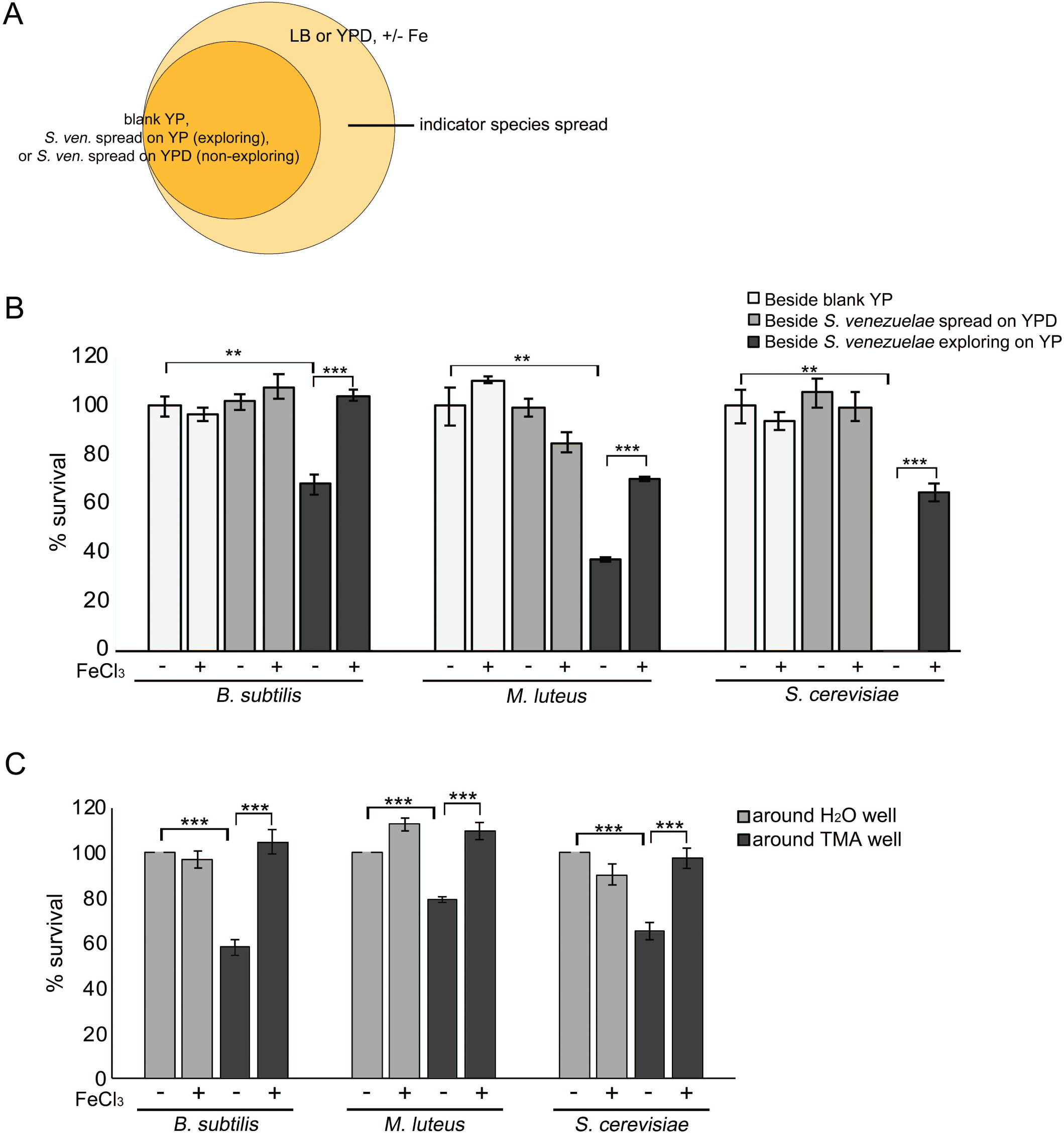
Iron supplementation restores the growth of microbes exposed to VOCs. **A.** Schematic of the experiment performed in **B**: Plates comprising uninoculated (blank) YP, *S. venezuelae* exploring on YP, or non-exploring *S. venezuelae* on YPD, were incubated in smaller dishes for 10 days. After 10 days, an indicator strain (*B. subtilis, M. luteus*, or *S. cerevisiae*) was spread on medium (+/-1 mM FeCl_3_ supplementation) in the outside dish. **B.** Quantification of *B. subtilis, M. luteus*, or *S. cerevisiae* colonies on medium with or without FeCl_3_ supplementation, following growth adjacent to exploring *S. venezuelae*, non-exploring *S. venezuelae*, or uninoculated medium **C.** Experiment conducted as in A and B, only with H_2_O or TMA solutions replacing YP/YPD inoculated with *S. venezuelae.* Plates were incubated at room temperature for 2 days. *B. subtilis, M. luteus*, or *S. cerevisiae* colonies were quantified following growth on medium with or without FeCl_3_ supplementation, adjacent to H_2_O or TMA solutions. For **B** and **C:** Values represent the mean ± standard error for three or four replicates, and statistical significance was determined using one-way analysis of variance (ANOVA), followed by Tukey’s multiple comparison test. Asterisks (*) indicate *p-*values (**: *p*-value of 0.01 to 0.005; ***: *p*-value below 0.005).

In contrast, when these microbes were grown next to exploring *S. venezuelae* on YP medium, colony numbers on the surrounding agar plates differed drastically depending on the iron supplementation status. Growing *B. subtilis, M. luteus* and *S. cerevisiae* adjacent to exploring *S. venezuelae* on medium without added iron led to a reduction in colony numbers by an average of 32%, 63%, and 100%, respectively, relative to controls (those grown on plates adjacent to blank YP or non-exploring *S. venezuelae* on YPD) (**Fig. 2B**). In each case, growth was partially (*M. luteus* and *S. cerevisiae*) or fully (*B. subtilis*) restored by iron supplementation (**Fig. 2B**). These data suggest that the alkaline environments created by exploring *S. venezuelae* reduced the viability of other soil microbes, at least in part by starving them of iron.

To determine whether this was a TMA-dependent phenomenon, or whether it was due to other volatiles produced by exploring *Streptomyces*, we set up equivalent assays where *S. venezuelae* colonies were replaced with water or TMA (11) (**Fig. 2C**). Around these water/TMA-containing wells were spread the three indicator organisms on agar medium, with or without iron supplementation. We quantified their growth after 2 days. For each microbe, the addition of iron had little impact on cell survival when cultures were grown next to water-containing wells (**Fig. 2C**): *B. subtilis* growth was unaffected, while there was a slight increase in growth observed for *M. luteus* and a slight decrease for *S. cerevisiae*.

In contrast, iron supplementation had a significant effect on survival and growth when cells were plated adjacent to TMA-containing wells. Exposure of *B. subtilis, M. luteus* and *S. cerevisiae* to TMA during growth on plates without iron supplementation led to reduced growth/survival of these strains by 42%, 21%, and 35% respectively (on average), compared with those grown on plates adjacent to water wells (**Fig. 2C**). As seen for the exploring culture experiments above, iron supplementation restored the growth of TMA-exposed cultures to levels equivalent to those around the water wells (**Fig. 2C**). This indicated that TMA emission by exploring *S. venezuelae* functioned to inhibit the growth of other microbes by limiting iron availability. It appears, however, that either exploring *S. venezuelae* produce more TMA than we were using in these assays, or additional volatile compounds also influence the growth of yeast, and to a lesser extent *M. luteus*, as the growth of these organisms was more strongly impacted by exploring *S. venezuelae*, than by TMA exposure.

### Desferrioxamines are produced during exploration

As iron supplementation could restore the growth of other microbes in the presence of TMA/*Streptomyces* volatile compounds, this suggested that these volatile compounds were responsible for creating a low-iron environment. This then raised the question of how exploring *S. venezuelae* dealt with these low iron conditions. To begin addressing this point, we examined the metabolic output of exploring cultures. Using liquid chromatography coupled with mass spectrometry (LC-MS), we compared the metabolites produced by exploring cultures (both *S. venezuelae* grown alone on YP medium and beside *S. cerevisiae* on YPD medium) versus non-exploring colonies (*S. venezuelae* grown alone on YPD or MYM medium) (**Fig. 3A-D**). We found that all exploring cultures produced analogs of the desferrioxamine or ferrioxamine (iron-complexed desferrioxamine) siderophore, including ferrioxamine B, D, and an aryl-functionalized ferrioxamine (**Fig. 3A-C**). Colonies exploring beside *S. cerevisiae* on YPD also produced the unusual ferrioxamine B+CH_2_, reported previously by Cruz-Morales *et al.* (36) (**Fig. 3D**). Importantly, ferrioxamines were not detected in non-exploring cultures, nor were they produced by *S. cerevisiae*, suggesting that siderophore production was specific to exploring *S. venezuelae* cultures under these growth conditions.

**Fig. 3.**
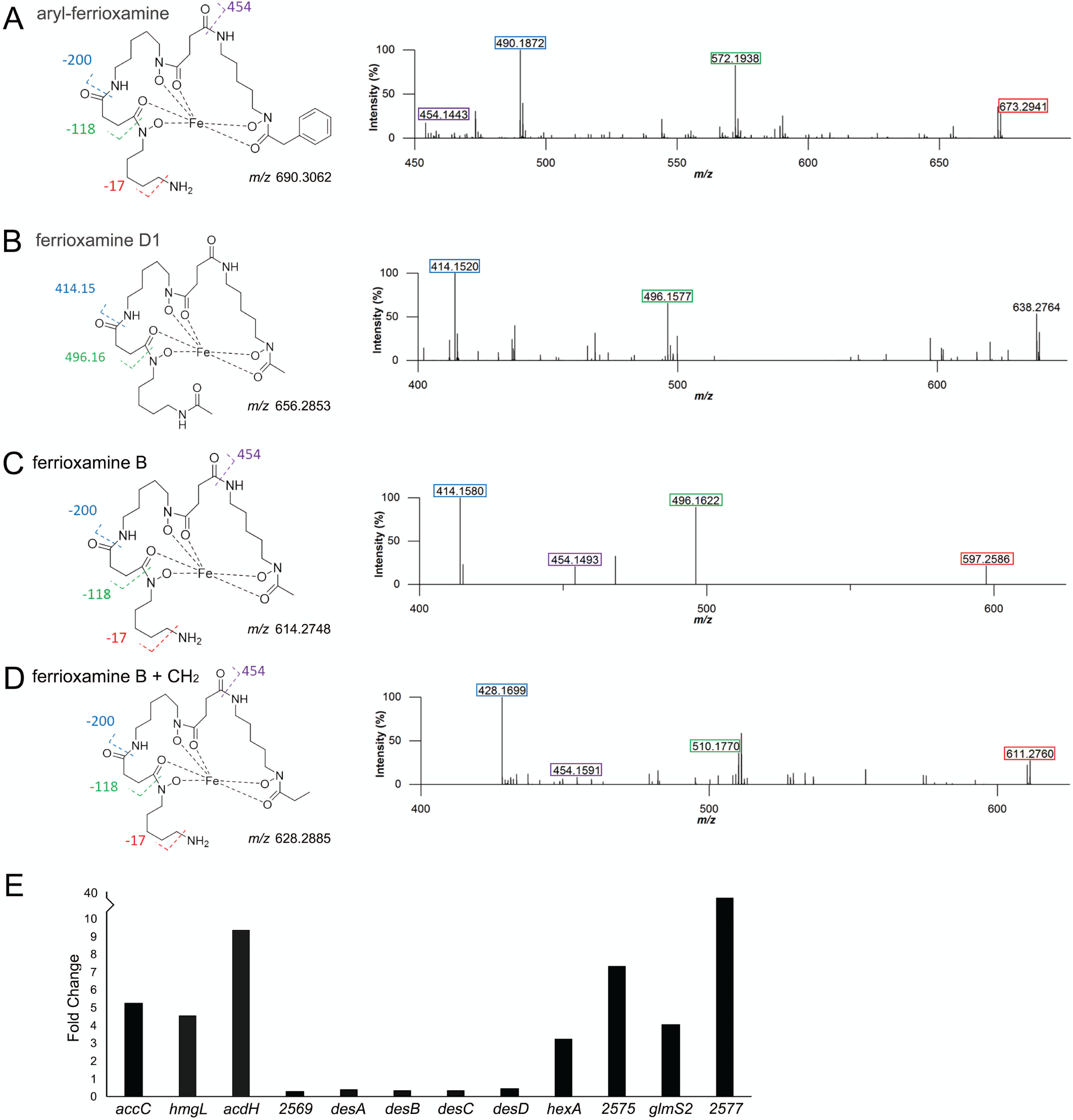
Explorer cells produce desferrioxamines. **A-D** show accurate mass fragmentation (MS^2^) data and molecular structures of associated ferrioxamine (iron-complexed desferrioxamine) compounds. No (des)ferrioxamines could be detected from non-exploring cultures (*S. venezuelae* spread on YPD). **A.** Aryl-ferrioxamine, from *S. venezuelae* exploring on YP and *S. venezuelae* exploring beside *S. cerevisiae* on YPD. **B.** Ferrioxamine D1, from *S. venezuelae* exploring on YP and *S. venezuelae* exploring beside *S. cerevisiae* on YPD. **C.** Ferrioxamine B, from *S. venezuelae* exploring on YP and *S. venezuelae* exploring beside *S. cerevisiae* on YPD. **D.** Ferrioxamine B + CH_2_ (where the CH_2_ is depicted as an ethyl group, as the methylene addition is on the right-hand side of the molecule and is most likely located in the acyl tail) from *S. venezuelae* exploring beside *S. cerevisiae* on YPD. **E.** Normalized transcript levels for genes in the *S. venezuelae* desferrioxamine biosynthetic cluster (as defined by antiSMASH (23)) in explorer cells, divided by those for non-exploring cells. The associated gene names or *sven* gene numbers are shown below each gene.

In parallel, we revisited our previously generated RNA-seq data (11), to determine whether exploring cultures were generally exhibiting a transcriptional profile consistent with iron starvation. While the regulation of iron homeostasis has not been studied in *S. venezuelae*, investigations in the closely related *Streptomyces coelicolor* have revealed that the desferrixoamine biosynthetic gene cluster (*desABCD*) is controlled by the iron-responsive regulator DesR/DmdR (20). DesR/DmdR binds to a site overlapping the -10 promoter region upstream of *desA*, repressing the expression of the *des* genes when iron is abundant. In iron-deficient conditions, transcription repression is alleviated, and transcription of the *des* operon increases as a result (20). It appears that a similar situation exists in *S. venezuelae*, which encodes a DesR/DmdR homologue (SVEN_4209) sharing 92% sequence identity (96% sequence similarity) with the *S. coelicolor* protein, and has a similar ‘iron box’ overlapping the promoter of the *des* operon (**Fig. S2**).

Unexpectedly, we found that *desABCD* (*sven_2570-73*) transcript levels were not upregulated in exploring cultures relative to static cultures (**Fig. 3E**). This suggested that exploring cultures were not exhibiting a traditional iron-starvation response. Furthermore, it implied that there must be some, as yet unknown, post-transcriptional regulatory mechanism functioning to enhance desferrioxamine production. It is worth noting that transcript levels for the genes flanking the *des* operon were significantly increased under exploring conditions (**Fig. 3E**), although it is not clear what role these gene products play in desferrioxamine synthesis.

### Oligopeptide transporters influence exploration and culture response to iron

Upon further analysis of our RNA-seq data, we observed that two of the most highly upregulated gene clusters in exploring cultures (*sven_5150-53* and *sven_4759-63*) encoded ATP-binding cassette (ABC) transporter systems (**Fig. 4A**). The closest characterized homologues to these genes in the model *S. coelicolor* system were in the *bldK* locus (*sco5112-16*). Recent work has suggested that BldK transporters function in ferrioxamine siderophore uptake (21). Increased expression for these two *bldK-*like transporter gene clusters in *S. venezuelae* implied that exploring cultures may adapt to alkaline, low-iron environments by coupling increased desferrioxamine production with enhanced ferrioxamine uptake.

**Fig. 4.**
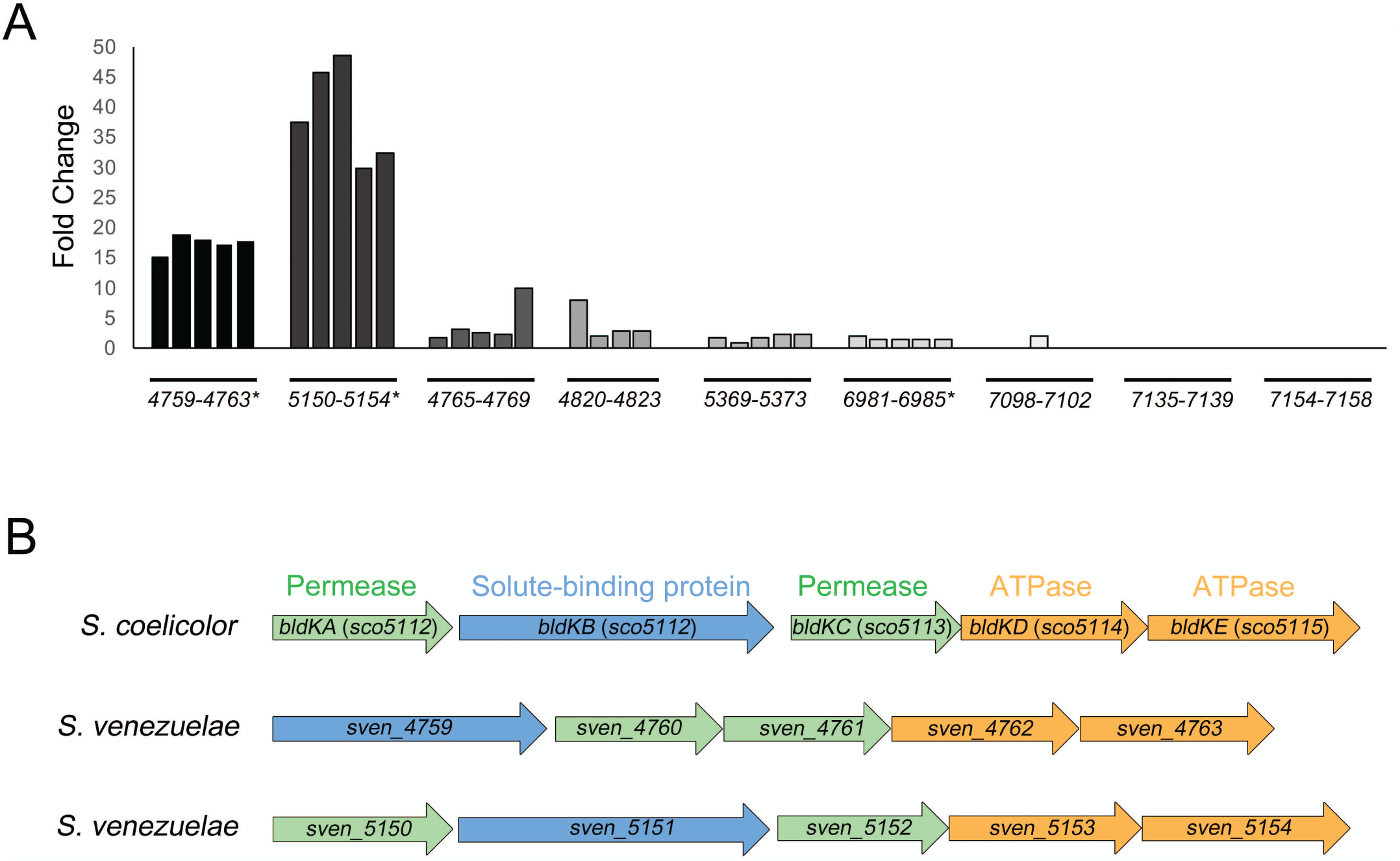
Upregulation of two gene clusters associated with siderophore transport in explorer cells. **A.** Normalized transcript levels for *S. venezuelae* clusters homologous to the *S. coelicolor bldKABCDE* locus, where normalized transcript levels in *S. venezuelae* explorer cells were divided by those in non-exploring cells. The associated *sven* gene numbers are shown below each cluster, and asterisks beside gene numbers indicate that there are genes in the cluster that are significantly differentially expressed in exploring versus non-exploring cultures, based on calculated *q*-values. **B.** Organization of the *S. coelicolor bldK* locus and the two *S. venezuelae bldK-*like clusters that are significantly and highly differentially expressed in exploring versus non-exploring cultures.

We searched for other genes in *S. venezuelae* that were homologous to the *S. coelicolor bldK* locus, and found *S. venezuelae* encoded seven additional *bldK-*like clusters (**Fig. 4A**). Of these, three were not expressed in exploring cultures or static cultures (*sven_7098-02, sven_7135-39, sven_7154-58*); three were expressed at low levels in both exploring and static cultures (*sven_4765-69, sven_4820-23, sven_5369-73*); and one was expressed at intermediate levels in exploring cultures, but was only modestly (∼1.7 fold) upregulated in exploring versus static cultures and is predicted to function in nickel transport (*sven_6981-85*) (22) (**Fig. 4A**). Thus, we focused our attention on the two highly expressed *sven_5150-53* and *sven_4759*-*63* clusters, in assessing their contributions to exploration and low iron adaptation.

To test whether the activity of these transporters affected exploration, we constructed mutations in key genes within each of the two upregulated *bldK-*like gene clusters (*sven_4759* and *sven_5151*). To account for the possibility of functional redundancy shared by these two transporters, we also created a double (*sven_4759/sven_5151*) mutant strain. We found that the single mutant strains each behaved like wild type on YP (exploration-promoting) medium (**Fig. S3**). However, the surface area of the double mutant was significantly larger (approximately three times greater) compared with wild type (**Fig. 5A,B**). This suggested that *sven_4759* and *sven_5151* were functionally redundant, and that their collective activity profoundly influenced exploration. The enhanced exploration capabilities observed for the double mutant could be reduced to wild type levels by complementing with a cosmid carrying an intact *sven_4759-63* operon (**Fig. S4**).

**Fig. 5.**
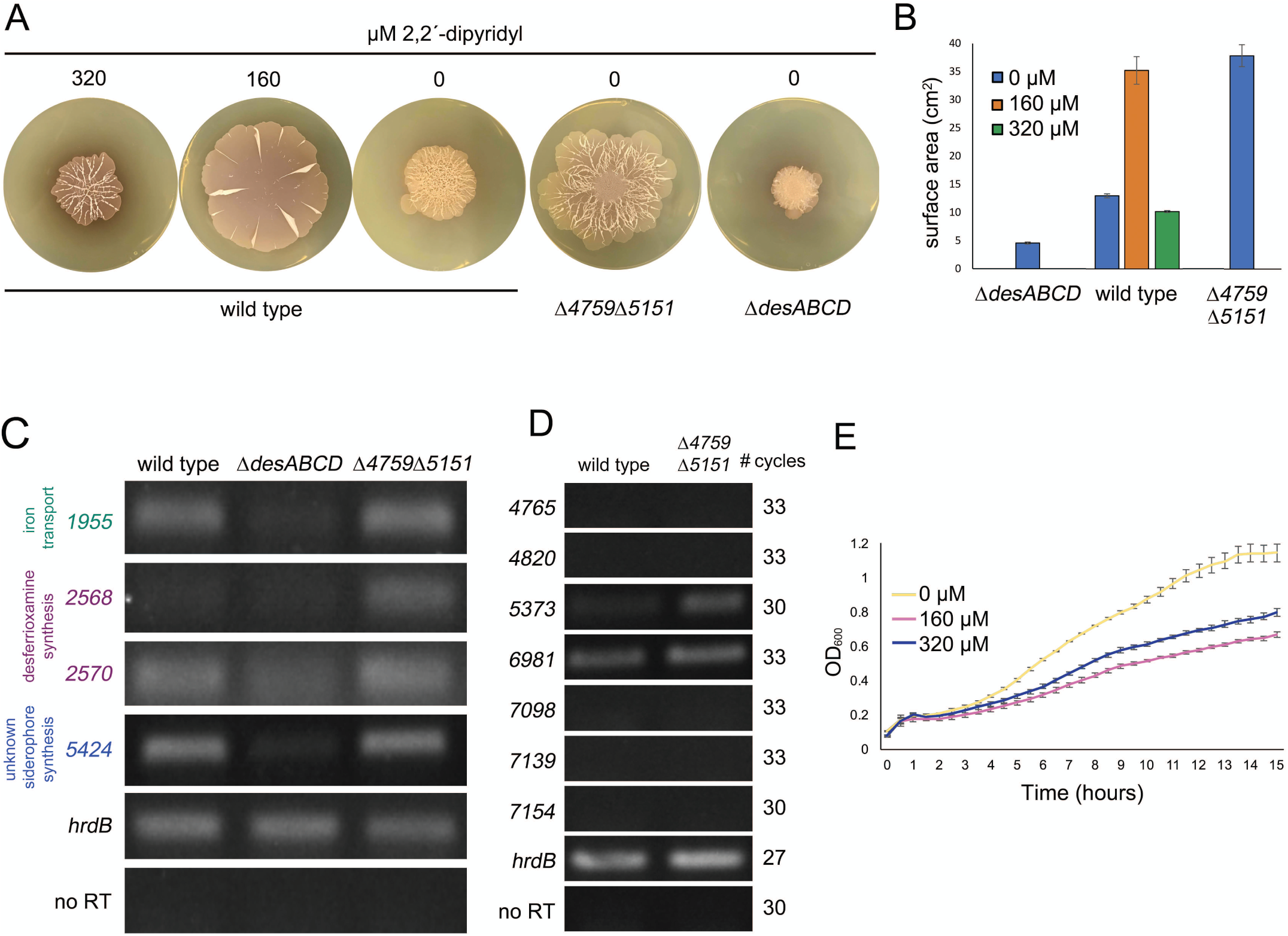
Iron uptake capabilities impact exploration. **A.** Wild type *S. venezuelae* exploring on YP with 0, 160 or 320 µM dipyridyl, alongside the *desABCD* mutant and the Δ*sven_4759*Δ*sven_5151* double mutant growing on YP agar medium. Plates were incubated for 10 days. Images are representative of three replicates per strain and dipyridyl concentration (where appropriate) **B.** Quantification of colony expansion by wild type *S. venezuelae*, the *desABCD* mutant, and the Δ*sven_4759*Δ*sven_5151* double mutant on YP agar, with 0-320 µM dipyridyl for wild type. Values represent the mean ± standard error for three replicates. **C and D.** Semi-quantitative RT-PCR using RNA isolated from wild type, Δ*desABCD* and Δ*sven_4759*Δ*sven_5151* double mutant strains grown for 8 days on YP medium. The vegetative sigma factor *hrdB* served as a positive control for RNA loading and RNA integrity, and no-RT reactions (using RNA as template with *hrdB*-specific primers) were included as negative controls to ensure a lack of DNA contamination of RNA samples and all PCR reagents. The number of PCR amplification cycles was optimized to ensure products were in the linear amplification range, and that no products were observed in the negative control for each reaction. For C, 30 cycles were conducted for all reactions apart from *hrdB*, where 27 cycles were used. Cycle numbers are shown to the left in panel D. Representative results are shown for experiments conducted in biological duplicate and technical triplicate. **E.** Growth curves of wild type *S. venezuelae* grown in liquid YP medium with 0-320 µM dipyridyl over 15 h. Values represent the mean ± standard error for three replicates.

These transporters were predicted to function in the uptake of iron-complexed siderophores, and their loss enhanced exploration. This led us to test whether low iron levels could generally enhance exploration. We grew wild type *S. venezuelae* on YP medium supplemented with increasing concentrations of the dipyridyl iron-chelator. On medium containing 160 µM dipyridyl, the surface area of exploring *S. venezuelae* was 2.7 times larger on average than that of colonies grown without chelator (**Fig. 5A,B**). This was consistent with the effect that we observed for the transporter mutants (on YP without dipyridyl), supporting our proposal that these transporters were involved in iron acquisition. Doubling the chelator concentration, however, resulted in wild type explorer colony sizes that were slightly smaller than seen in the absence of chelator, suggesting that there was a threshold level of iron required to promote robust exploration.

These observations prompted us to create a desferrioxamine mutant strain (Δ*desABCD*). We expected to see increased exploration for this mutant, as we had for the transporter mutant. Instead, we observed reduced exploration for the *des* mutant relative to wild type (**Fig. 5A,B**), suggesting that iron uptake by this strain may fall below the threshold needed for efficient exploration.

In an effort to reconcile the conflicting phenotypic observations for the transporter and *desABCD* mutant strains, we wanted to determine whether the expression of other siderophore clusters or transporters were altered in these strains, relative to each other and to wild type. We isolated RNA from each strain after 8 days of exploration, and analyzed the expression profiles of genes predicted to direct siderophore production and uptake in these strains using semi-quantitative RT-PCR. *S. venezuelae* encodes five predicted siderophore biosynthetic clusters (as predicted by antiSMASH, see **Table 1**) (23). Transcript levels associated with genes in the desferrioxamine biosynthetic cluster (*sven_2568* and *sven_2470/desA*) were higher in the double transporter mutant compared to wild type (**Fig. 5C**), consistent with what has been observed previously for *bldK* mutants in *S. coelicolor* (21). In contrast, genes in the other four siderophore-encoding biosynthetic clusters were unaltered in wild type and transporter mutant strains (**Fig. S5**). Interestingly, the *desABCD* mutant exhibited reduced expression of several other transporter genes and siderophore biosynthetic clusters when compared with wild type and the transporter mutant strains (**Fig. 5C**). This altered expression may serve to further magnify the iron uptake defects of this strain, and explain its reduced capacity to explore.

We also examined the expression of the other *bldK*-like genes in the double transporter mutant strain, to determine whether any of these might be upregulated, such that their products could compensate for the loss of the two transporters. We found that transcription of genes in most clusters was undetectable, with the exception of the *sven_6981-*containing cluster, which was unchanged between wild type and mutant, and the *sven_5373*-containing cluster, whose expression was upregulated in the double mutant background (**Fig. 5D**). We speculate that the transporter encoded by this cluster may facilitate low-level ferrioxamine-uptake in the absence of the two primary transporters, ensuring sufficient intracellular iron levels for robust exploration to occur.

Finally, we sought to understand more about the mechanism underlying the increased exploration colony size observed for the siderophore transporter mutant and the wild type strain growing under low iron conditions. We reasoned that the larger surface area observed for these strains, relative to wild type grown without chelator, could be due to enhanced exploration, or it could be a result of increased growth. To differentiate between these possibilities, we grew wild type *S. venezuelae* for 15 hours in YP liquid medium, with or without the dipyridyl chelator. We found that chelator supplementation led to generally reduced *S. venezuelae* growth (**Fig. 5E**). This suggested that the enhanced exploration observed for the wild type strain growing on 160 µM dipyridyl, as well as the transporter mutant strain, was most likely the result of increased colony expansion, and not more rapid growth.

### Low iron environments can be created by interspecies interactions

*Streptomyces* live alongside many other bacteria and fungi in the soil where there is intense competition for key nutrients, including iron. Given that low iron both reduced the survival of other microbes and enhanced exploration, we wondered how microbial competition for iron would impact *S. venezuelae* exploration. We focused our attention on interactions between *Streptomyces* and *Amycolatopsis* bacteria. These bacteria have been isolated from the same soil samples, suggesting that they may well interact in the environment (24). Furthermore, previous work has demonstrated that *Amycolatopsis sp.* AA4 can pirate desferrioxamine E from *S. coelicolor* in low iron environments, with this iron sequestration also inhibiting the aerial development of *S. coelicolor* (24, 25). We recapitulated this experiment using *Amycolatopsis orientalis* sp. *lurida* and observed the same defect in aerial development for *S. coelicolor*, suggesting that this species can also sequester *Streptomyces* siderophores (**Fig. S6**).

We next tested whether such siderophore piracy could affect *S. venezuelae* exploration. We grew *S. venezuelae* beside *A. orientalis*, as described above for *S. coelicolor*, on either YP agar or YP supplemented with dipyridyl. Following a 7 day incubation, we found the surface area of *S. venezuelae* had increased by >100% when grown beside *A. orientalis* on YP agar without chelator (**Fig. 6**). This was analogous to the growth of *S. venezuelae* alone on YP medium supplemented with moderate concentrations (160 µM) of chelator (**Fig. 6**). Interestingly, combining proximal *A. orientalis* with further iron sequestration (through the addition of 160 or 320 µM dipyridyl) led to a decrease (14% and 76%, respectively) in the surface area of exploring *S. venezuelae*, suggesting that iron levels under these conditions were below the threshold required for optimal exploration.

**Fig. 6.**
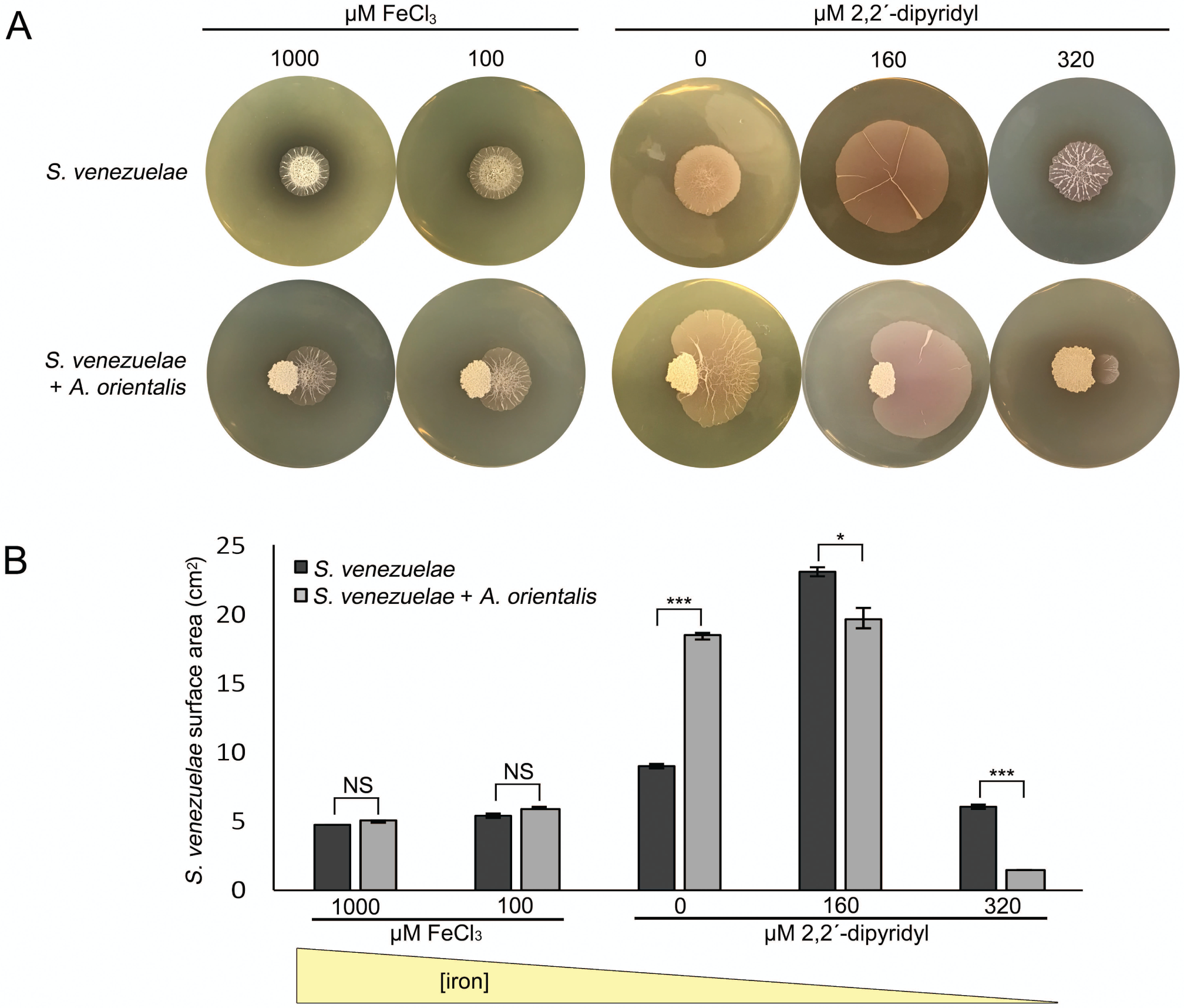
New interspecies interactions alter exploration by creating low iron environments. **A.** *S. venezuelae* grown alone (top row) or beside *A. orientalis* sp. *lurida* on YP supplemented with FeCl_3_ or dipyridyl. Plates were incubated for 7 days, and all images are representative of three replicates per condition. **B.** Quantification of *S. venezuelae* grown alone or beside *A. orientalis* on YP agar with decreasing levels of available iron (left to right). All values represent the mean ± standard error for three or four replicates. Asterisks indicate statistically significant differences (*: *p*-value 0.05 to 0.01; ***: *p*-values below 0.005, NS: no significant difference) as determined by a student’s t-test.

Our results suggested that *A. orientalis* promoted increased *S. venezuelae* exploration in the same way as low iron growth conditions. To test this hypothesis, we inoculated *S. venezuelae* beside *A. orientalis* on YP medium supplemented with increasing concentrations of FeCl_3_ for 7 days (**Fig. 6**). For the different FeCl_3_ concentrations tested, the surface area of *S. venezuelae* was nearly identical when grown alone or beside *A. orientalis.* This indicated that the exploration-promoting effects of *A. orientalis* could be suppressed by excess iron, implying that iron sequestration by *A. orientalis* was responsible for enhancing *S. venezuelae* exploration.

### Glucose trumps iron in the hierarchy of nutrients affecting exploration

Our results to this point suggested that low iron levels, irrespective of whether this was due to VOC-mediated alkalinization, iron uptake defects, or siderophore sequestration, resulted in enhanced exploration. This led us to question whether low iron could overcome the exploration-inhibitory effects observed for other nutrients. Previous work revealed that exploration was glucose-repressible (11), and thus we wanted to determine whether low iron conditions could alleviate the need for a low-glucose environment. We grew *S. venezuelae* and the Δ*sven_4759*Δ*sven_5151* double mutant strain on YP plus glucose (YPD) plates supplemented with 160 µM or 320 µM of the dipyridyl iron chelator for 10 days. We found that *S. venezuelae* failed to explore under these conditions (**Fig. S7**). This indicated that the presence of glucose was sufficient to override the effects of iron deficiency, with respect to exploration promotion.

We also tested whether additional iron could inhibit exploration. We grew wild type *S. venezuelae* on YP medium (exploration-promoting medium) and MYM medium (classic development-promoting medium) with increasing concentrations of iron (**Fig. 7**). *S. venezuelae* explored on YP with iron levels ranging from 0-10 mM, although the surface area was reduced slightly at the highest iron levels when compared to strains grown without added iron. Notably, *S. venezuelae* failed to grow on MYM medium when iron levels exceeded 2.5 mM, but exploring cultures remained viable when growing on concentrations at least four times that. This raised the interesting possibility that exploration could also protect *S. venezuelae* from otherwise toxic levels of iron in their environment by reducing the amount of bioavailable iron through environmental alkalinization.

**Fig. 7.**
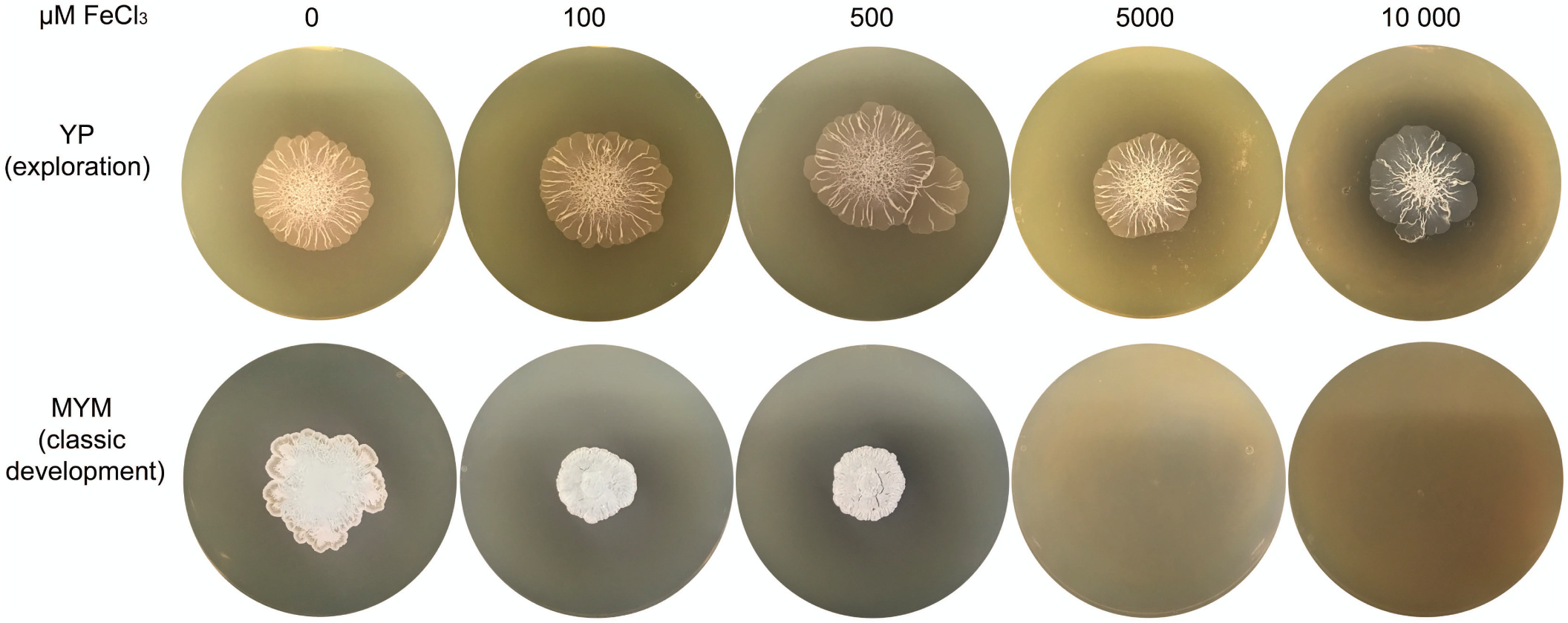
*S. venezuelae* growth on medium supplemented with iron. Wild type *S. venezuelae* grown on YP medium (top row; exploration-promoting) or MYM medium (bottom row; classic growth medium) supplemented with 0-10,000 µM FeCl3. Images are representative of three replicates per media type, per iron concentration.

## DISCUSSION

Our work here reveals a new role for bacterial VOCs in modulating nutrient availability and microbial community behaviour, and expands the repertoire of interspecies interactions that can impact exploration. We show the release of TMA by *Streptomyces* explorer cells dramatically alters the pH of their surrounding environment, and in doing so, reduces the survival of nearby soil microbes by starving them of iron. Within these iron-depleted niches, exploration is enhanced and *Streptomyces* ensure maximal iron uptake by secreting siderophores, and upregulating genes encoding putative desferrioxamine transport systems. Collectively, our results reveal exploration to be not only an effective mechanism for dealing with competition for iron and iron toxicity, but also a potent mediator of nutrient availability in the environment.

### Volatile compounds impact microbial community dynamics by controlling nutrient availability

*Streptomyces* exploration is coordinated by the volatile molecule TMA, whose release raises the pH of the surrounding environment (11). Remarkably, TMA functions as both a communication signal, inducing other *Streptomyces* to explore, and as a competitive weapon, reducing the survival of other microbes. The antibacterial and antifungal properties of TMA appear to be tied to its nutrient modulatory effects. Alkaline conditions create an inhospitable environment for many microbes by reducing the levels of bioavailable iron (17, 18, 26). We show that iron supplementation can restore the growth of otherwise susceptible bacteria and fungi in the presence of TMA, suggesting that iron starvation is at the heart of the TMA antimicrobial effects.

Microbes can modify their environment through nutrient depletion, or metabolite excretion, and in doing so, can impact the growth and dynamics of their surrounding community members. Increasingly, volatile compounds are being recognized as important contributors to soil nutrient status. For example, microbial volatiles can generally impact carbon dynamics in the soil (27). Species-specific volatiles have also been shown to enhance the availability of reduced sulfur, which in turn promotes the growth of sulfur-deficient plants (28). We can now add volatile-induced iron starvation through increased environmental pH to the effects ascribed to microbial volatiles.

### Promoting exploration in iron-depleted environments

*Streptomyces* exploration is enhanced during growth in low iron environments, although below a certain level, exploration stimulation ceases. Iron is limited in the soil (24, 29). Exploratory growth in iron-limited environments could allow *Streptomyces* to access nutrients in more distant locations. We have determined that exploration is both rapid and remarkably processive; explorer cells spread over surfaces at a rate that is ∼12 times faster than has been seen previously for *Streptomyces* colonies (11), and it is not ye t known how exploration is stopped. Our data suggest the relentless nature of exploration could be explained by a positive feedback loop, with iron acting as a central player: explorer cells produce TMA, TMA emission leads to increased pH and reduced iron availability, explorer cells spread more to get more iron, TMA continues to be produced, and the cycle continues (**Fig. 8**). Indeed, a recent study modeling the effects of pH on bacterial growth demonstrated that when environmental modification is beneficial for a bacterium, there is a positive feedback on their growth: the more they change the environmental pH, the more cells grow, and the more they can continue to alter pH (26).

**Fig. 8.**
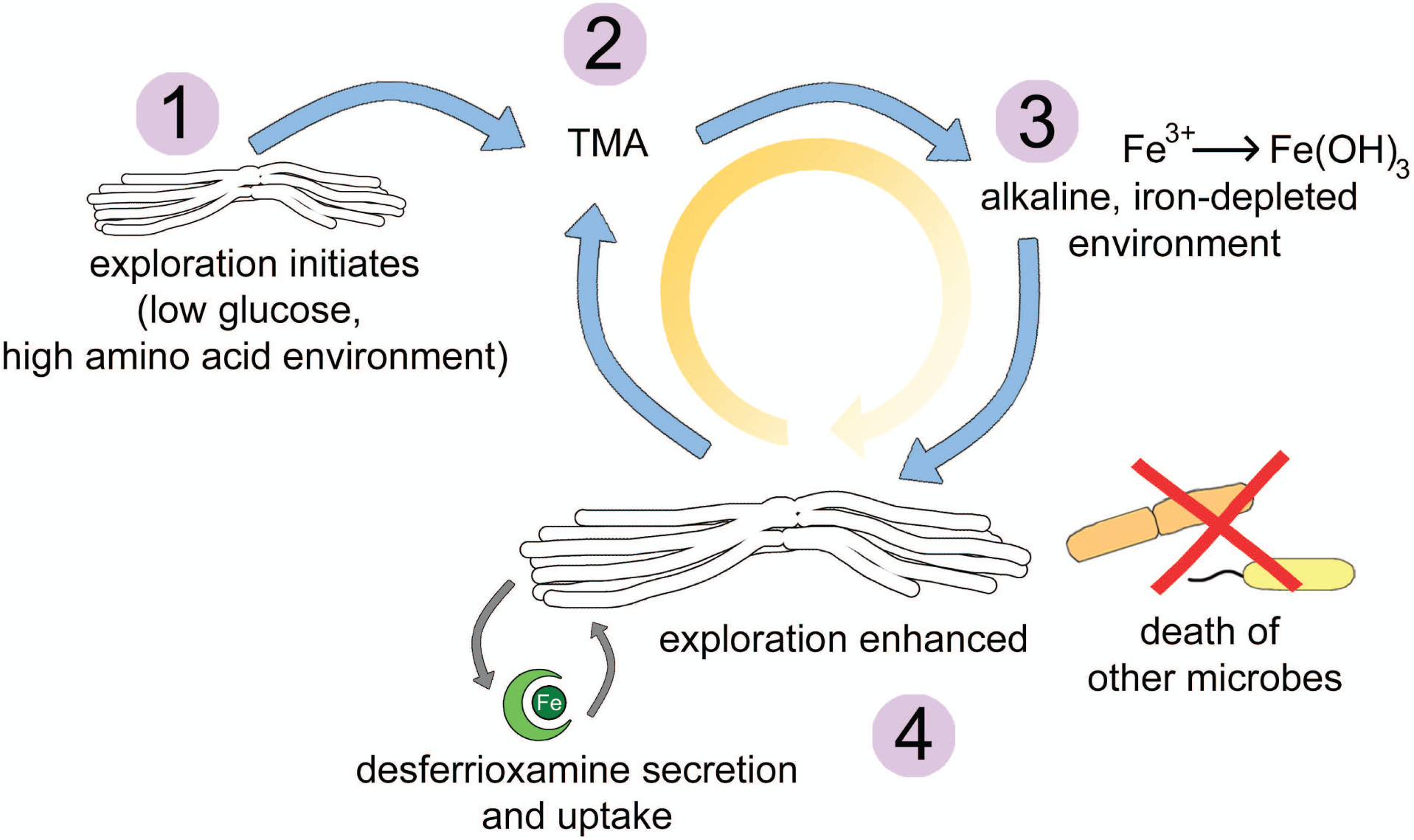
*S. venezuelae* explorer cells thrive and kill other microbes in alkaline, low-iron environments. 1. *S. venezuelae* exploration is triggered by a combination of low glucose and high amino acid concentrations. 2. Explorer cells release the VOC TMA into the surrounding environment. 3. TMA raises the pH of the environment, concomitantly reducing the solubility and bioavailability of iron. 4. To cope with low iron conditions, *S. venezuelae* explorer cells release desferrioxamines. These siderophores return solubilized iron to the cells. At the same time, *S. venezuelae* exploration is enhanced, perhaps as a mechanism to reach more iron-rich environments. This enhancement of exploration leads to increased TMA production, creating a positive feedback loop (orange arrow): TMA depletes iron, explorer cells spread to get more iron, TMA production continues, and the cycle repeats. Within these alkaline, iron-depleted environments, the growth of other bacteria and fungi is reduced.

Within the exploration feedback cycle, several additional factors could account for the increase in exploration surface area in low iron environments. The classic *Streptomyces* life cycle involves a transition from vegetative hyphal growth to raising reproductive aerial structures. For *S. coelicolor*, the addition of iron chelators to the growth medium prevents colonies from raising aerial hyphae or sporulating, and instead locks cells in the vegetative growth phase (24). Explorer cells share many properties with vegetative hyphae (11), and it is possible that reduced iron availability helps to inhibit aerial development, thereby enhancing the rate of exploration by preventing entry into the classical reproductive differentiation phase.

We also noted that colonies growing on YP supplemented with the dipyridyl chelator, or in association with *A. orientalis*, were much flatter and less structured than colonies growing alone on YP. Other bacteria (*e.g. P. aeruginosa*), require iron for biofilm assembly: low iron triggers increased surfactant production and motility, which in turn reduces biofilm structure (30, 31). Similar connections between iron availability and motility have been made in other microbes (32–35). *Streptomyces* exploration is phenotypically similar to sliding motility – a form of passive motility driven by growth and facilitated by the release of a surfactant. It is possible that low iron enhances surfactant production by explorer cells. This in turn could alter colony architecture and promote greater motility, while at the same time increase the colony’s ability to scavenge for iron.

How *Streptomyces* sense low iron levels during exploration remains unexplained. It does not appear to be mediated by the major iron repressor DesR/DmdR, as expression of the *des* cluster, a known regulon member in other streptomycetes, is unaltered during exploration. It is also not clear what controls the expression of the two transporter-encoding clusters, as these do not have an ‘iron box’ present in their promoter regions. It will be interesting to determine whether this response is coupled with pH-sensing, given the profound effect of high pH on iron availability.

### *Interspecies effects on* Streptomyces *metabolism and exploration*

*Streptomyces* exploration was initially discovered as a response to co-incubation with the yeast *S. cerevisiae*. Specifically, *S. cerevisiae* served to deplete glucose from the medium, and it was this change in carbon source availability that promoted the onset of exploration. We show here that other organisms capable of ‘stealing’ siderophores, such as *Amycolatopsis*, can also influence the rate of exploration. Intriguingly, *S. venezuelae* secretes a suite of differentially modified siderophores during exploration, including unusual molecules for which there is only one report in the literature (36). It will be interesting to determine whether some of these molecules are more amenable for uptake by *S. venezuelae* than by other organisms, and whether these modified compounds provide resistance against siderophore cheaters. Alternatively, the high pH environment may promote differential siderophore tailoring, as has been noted for marine microbes that produce more amphiphilic ferrioxamines (37). In exploring *S. venezuelae*, this maybe the result of increased expression of the genes flanking the core *desABCD* operon, and/or the effect of modulating the availability of different precursors.

Collectively, our work reveals that exploring *Streptomyces* colonies can alter nutrient availability thorough volatile compound emission, and that this activates a positive feedback cycle that promotes continued exploration. This nutrient modulation further drives changes in the dynamics and composition of the local microbial community.

## Supplementary Figure Captions

**Fig. S1. Exploring *S. venezuelae* creates an alkaline environment through the release of volatile compounds**. A smaller dish containing YP medium (left) or YP inoculated with *S. venezuelae* (right) was incubated within a larger dish containing LB medium. After 10 days, when *Streptomyces venezuelae* exploration was well-underway, bromothymol blue pH indicator dye was spread on the larger agar. Blue indicates VOC-induced alkalinity.

**Fig. S2. Organization of the desferrioxamine biosynthetic cluster and its control by DesR/DmdR. A.** The desferrioxamine biosynthetic clusters, with gene names and associated *sven* numbers shown below each gene. Promoter locations are shown as black arrows., and colours indicate predicted gene function. Promoter locations are shown as black arrows. Promoter region upstream of *desA* is indicated with a box in the cluster diagram, and the sequence is shown below with the predicted DesR/DmdR binding site highlighted in grey. **B.** Alignment of the promoter regions for *desA* in *S. coelicolor* (top) and *S. venezuelae* (bottom). **C.** Alignment of the DesR/DmdR regulators from *S. coelicolor* (SCO4394) and *S. venezuelae* (SVEN_4209).

**Fig. S3. Mutations within individual *bldK-*like gene clusters do not impact exploration.** Wild type *S. venezuelae*, Δ*sven_4759* and Δ*sven_5151* grown on YP medium. Plates were incubated for 10 days. Images are representative of three replicates per strain per condition.

**Fig. S4. Complementation of the double *bldK* mutant.** Wild type *S. venezuelae*, the double *bldK* mutant, and the double *bldK* mutant complemented with a cosmid carrying the wild type sequence of *sven_4759* were grown on YP medium for 10 days.

**Fig. S5. Expression profiles of genes involved in siderophore synthesis and iron uptake.** Semi-quantitative RT-PCR using RNA isolated from wild type, *desABCD* mutant, and Δ*sven_4759*Δ*sven_5151* double mutant strains grown for 8 days on YP medium. The vegetative sigma factor *hrdB* served as a positive control for RNA loading and RNA integrity, and no-RT reactions (using RNA as template with *hrdB*-specific primers) were included as negative controls to ensure a lack of DNA contamination of RNA samples and all PCR reagents. Representative results are shown for experiments conducted in biological duplicate and technical triplicate.

**Fig. S6. *Amycolatopsis* inhibits aerial hyphae formation by *S. coelicolor*.** Left: *S. coelicolor* grown alone. White colony areas are aerial hyphae. Right: *S. coelicolor* grown beside *Amycolatopsis orientalis* sp. *lurida.* Shown are two replicates per plate, with both plates having been incubated for 14 days.

**Fig. S7. Glucose represses exploration, even in the presence of low iron.** Wild type *S. venezuelae* and the Δ*sven_5759*Δ*sven_5151* mutant strain grown on 0 or 160 µM 2,2′-dipyridyl. Two replicates of each strain were spotted on each plate, and strains were incubated for 10 days.

